# Metabolic ecology, microbial community structure, and gene-centric metagenomics

**DOI:** 10.1101/2025.08.05.668563

**Authors:** José Ignacio Arroyo, Beatriz Díez, Pablo A. Marquet

**Author notes:** Corresponding authors: José Ignacio Arroyo, Pablo Marquet. José Ignacio Arroyo, Santa Fe Institute, 1399 Hyde Park Road, Santa Fe, NM 87501, USA. Beatriz Díez, Universidad Mayor, Santiago, Chile.

## Abstract

Metabolic ecology includes predictions for biological rates from molecules to ecosystems, but despite the broad range of its scope, its applicability to metagenomes remains an open question. Here, we integrate metabolic ecology with principles of DNA shotgun sequencing to generate specific testable predictions for the metagenomic structure of microbial communities. We start by predicting a scaling relationship between population and assemblage abundance with genome size. This allows us to simplify DNA shotgun sequencing equations for the reads of a population and a gene. Then we derive the temperature dependence of population and assemblage abundance using the volume-temperature and abundance-volume rules and integrate them with simplified DNA sequencing equations to show that these predictions are compatible with a metagenomic framework. In addition, we derive predictions for the temperature dependence of the structure (abundance and richness) of genes and groups of related genes (e.g., metabolic pathways). To test our model, we provide some example data from human, aquatic, and terrestrial microbiomes from recent global projects. All predictions were supported by the observed data. Our model, derived from the integration of first principles, provides a mechanistic basis for variation in the structure of microbial community genomes in environmental and host-associated ecosystems.

## Introduction

Theories allow scientists to make predictions, anticipate phenomena, and recommend policy based on scientific evidence (Carnap 1946, Scriven 1959, Scheiner and Willig 2008, Marquet et al. 2014). Among theories, of special importance are those that are not only empirical approximations (i.e., provide good fits to data), but are derived from first principles, and thus provide a solid mechanistic basis to explain the phenomena under study and a logical interpretation of the parameters they contains. It has been argued that an efficient theory should be a relatively simple mathematical model (i.e., with few free parameters) derived from first principles that make testable predictions of wide applicability (Marquet et al. 2014). Within the definition of an efficient theory, only a few ecological theories can be counted, including life history, optimal foraging, and metabolic theory (of ecology; MTE), among others (Marquet et al. 2014).

The MTE attempts to explain the rates and patterns of the transformation of matter and energy in biological systems based on a few fundamental variables: body size and temperature. This framework emerged from the integration of two well-known biological relationships: Kleiber’s law relating body size and metabolic rate, and Arrhenius’ equation, relating temperature and (bio)chemical reaction rates (Brown et al. 2004). Since metabolic rate is the sum of biochemical reactions within a living entity, it would also be determined by the Arrhenius equation; thus, both effects were united to explain metabolic rate. As originally formulated, the MTE equation is based on first principles. Although the metabolic rate-mass relation was originally an empirical pattern, West et al. (1997) derived a mechanistic model to explain this scaling based on the constraints that fractal distribution networks, in plants and animals, impose on metabolisms, whereas in unicellular organisms, diffusion of metabolites seems to be more important (Gallet et al. 2017). Similarly, the Arrhenius equation, although justified in physical terms, is mostly empirical, and although it provides a good fit to empirical data, it fails to account for the common nonlinearities underlying reaction kinetics and temperature performance curves (see Delong et al. 2017). Any theory is susceptible to improvement. To have an equation derived from first principles but simple we have proposed a general model of temperature dependence derived from the fundamental (Eyring-Polanyi) equation of the transition state theory, which is currently the most accepted model for chemical kinetics (Zhou 2010), provides a good representation of temperature response kinetics including nonlinearities, and is rooted in the first principles of thermodynamics. The Eyring-Polanyi equation is

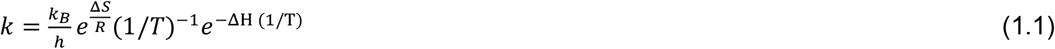

Here *k* is a (first order) reaction rate (s^−1^), h is the Planck constant (J/s), *ΔS* is the entropy change (J K^−1^), and *ΔH* is the enthalpy change (J). If replacing *ΔS* in terms of heat capacity, we obtain a new model which has the form of a power law with an exponential cut-off (Arroyo et al. 2022),

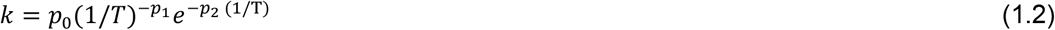

where 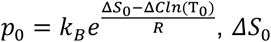 is entropy at *T*_0_, *T*_0_ is a temperature of reference (commonly 25°C, i.e., 298.15 K), *ΔC* is heat capacity (the rate of change of entropy) (J K^−1^), *T* is temperature in Kelvin, *R* is gas constant (J K^−1^ mol^−1^), 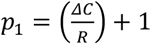and *p*_2_ = *ΔH*/*R, ΔH* is enthalpy change (energy transferred to the environment) (J). Negative values of *ΔC* gives an asymmetric concave curve that accounts for the breakpoint at optimum temperature, characteristic of temperature response curves. Notice that Eq.1.2 becomes Eq. 1.1 for *p*_1_ = 1, then Eq. 1.1 is nested within Eq. 1.2.

Predictions in metabolic ecology have included many variables from molecules to ecosystems, including population carrying capacity, species richness, among others (Brown et al. 2004, West et al. 2005, Sibly et al. 2012). Despite all these advances, there is still potential for this theory to be expanded due to temperature being pervasive. Here we visualize the opportunity to integrate metabolic ecology with metagenomic principles to predict variation in population and community genomes. Metagenomics, or the study of ecological communities through the sequencing of all the environmental DNA, has proven to be an invaluable tool to characterize the complexity of microbial communities (Segata et al. 2013, Escobar-Zepeda et al. 2015, Aguiar-Pulido et al. 2016). A metagenomic study includes the extraction of all the environmental DNA of a given sample that could be environmental or host-associated, its sequencing leading to a set of reads (i.e., a short DNA sequence of different length depending on the technology used). These reads can be studied directly in order to identify to which gene and taxa are more similar to (a gene-centric approach). The assignment to a gene and taxa is carried out through sequence alignment of reads against a database of characterized sequences whose taxonomic and/or gene affiliation is known. Databases typically include algorithms to assign unknown sequences to well-known groups of genes that typically correspond to ‘trees’ (e.g., a gene family), ‘paths’ (a metabolic pathway), or ‘networks’ (a metabolic network or protein structure) of genes. For a single sample, after this functional and taxonomic characterization of reads, the output will be a vector of the number of reads assigned to each functional category and a vector of the number of reads assigned to each taxon. In comparative metagenomic studies, instead of a vector, the output will be a matrix of *n* samples by *m* functional categories or *p* taxonomic groups (species, genus or any level defined by the investigator). Eventually, the relative or normalized abundance of reads can be compared among samples to assess the effects of different environmental factors on the relative abundance of a given function or taxa. This gene-centric comparative metagenomic approach has allowed to characterize of the structure (meaning the abundance and richness of taxa and their functions) of microbiomes, including human (Huttenhower et al. 2012), ocean (Sunagawa et al. 2012), and topsoil (Thompson et al. 2017), among others. These studies have shown that the abundance and richness of taxa and their functions, i.e., commonly referred as the structure of the microbiome, vary with environmental variables. For instance, the human microbiome varies according to organ type (Costello et al. 2009), whereas in the global ocean and topsoil, changes in microbiome structure are driven by temperature and pH, respectively (Sunagawa et al. 2015, Fierer 2017). Another approach in metagenomics, genome-centric metagenomics, includes the assembly of reads into contigs (set of overlapping reads that form a unit) and its incorporation into genome bins (sets of contigs that form a circular DNA sequence), allowing the quantification of variation of a population in one or several samples (Sharon and Banfield 2013) and ultimately the reconstruction of their genomes and metabolomes (Lawson et al. 2017) and predictions of their growth rates (Korem et al. 2015). Gene-centric metagenomics has evolved from descriptive to comparative and ultimately statistical and systems biology approaches (Wooley et al. 2010, Thomas et al. 2012, Quince et al. 2017). Despite the progress in metagenomics only a few models have been derived to predict the structure of community genomes (for instance based on biogeochemical models; Reed et al. 2014) and to the best of our knowledge, a model for the temperature dependence of the structure (relative abundance and richness) of taxa and their genes has not been derived. Here we integrate metabolic ecology and DNA sequencing theory to deduce the temperature dependence of the structure of metagenomes.

### Theoretical predictions

Predictions in metabolic ecology are made for units of body mass or area of a community or ecosystem, but commonly, microorganisms are quantified in distinct units. Their body size (i.e., cell) is measured in units of volume, and the individuals or richness of a community is also measured in units of volume, although using metagenomics, it is measured in units of individuals. To reconcile these different units in our derivations, we assume that i) earlier predictions for body size (kg) are also valid for unicellular cell volume (*μm*^3^) and ii) those made for unit area are valid for unit volume, and iii) Moreover, we assume that the number of individuals scales with volume; therefore, predictions would also be valid for communities measured by unit individuals. These three assumptions are supported (see, for example, Schmid et al. 2000).

### Simplifying DNA sequencing theory using ecological rules

We will derive our framework in three steps. First, we will derive scaling relationships between abundance and genome size (at the level of population and community) to subsequently simplify metagenomic sequencing theory. Second, we will use ecological principles to predict taxonomic community structure characterized using metagenomics, and third, we will use another set of ecological principles to predict the functional (genes and metabolisms) community structure.

We will start combining the cell size-genome size and population abundance-cell size relationship to derive relationships between abundance and genome size. The relationship between cell size and genome size has been shown in prokaryotes and unicellular eukaryotes (Hjort and Bernander 1999, Cavalier-Smith 2005, West and Brown 2005, Connolly et al. 2008, Olefeld et al. 2018), and the mechanism for its origin has been hypothesized to be related to optimization of rates leading to physical constraints (West et al. 1999, Gallet et al. 2007, Cavalier-Smith 2005). Particularly, the mechanism could be different for prokaryotes and eukaryotes, and for this last group, the debate has a long history (Gallet et al. 2007, Cavalier-Smith 2005). The cell volume (*μm*^3^)-genome mass (picograms, pg) relationship can be approximated as a power law

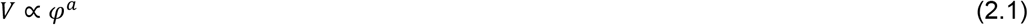

We can derive an alternative to Eq. 2.1 by taking averages (Savage 2004)

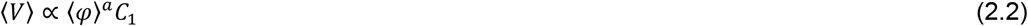

Where *C*_1_is a constant (see Savage 2004).

We will combine the cell volume-genome mass pattern with the relationship between population abundance and body size, which has been commonly documented in (at least) four different forms (White et al. 2007). For instance, at the global scale, mean population abundance decreases with mean size as reported by Damuth (1981, see also Belgrano et al. 2002) with a slope of −3/4, leading to the hypothesis that population energy use is invariant with mass, the energetic equivalence rule (EER, Damuth 1981, Blackburn and Gaston 1999). Another form of the abundance-size relationship is the assemblage abundance and mean cell volume of the assemblage (White et al. 2007, see our eq. 3.2). At the local scale, the relationship between population abundance and body size or volume (eq. 3.1) has been shown to be valid in local communities (Marquet et al. 1990, Schmid et al. 2000, Marquet et al. 2005). In general, the theory predicts that these relationships should apply to the maximum density that a species can achieve as a function of its size (Enquist et al. 1998, Belgrano et al. 2002). In microbial communities, abundance is commonly measured per unit of volume and size as cell volume, then the relationship is abundance-volume (Christensen et al. 1999, Cermeno et al. 2006, Sjostedt et al. 2012, Polyanskaya et al. 2015). Thus, the volume (V) scaling of local population abundance (*N*), assemblage abundance (*N*_T_) and global mean population abundance ⟨*N*⟩ can be approximated as

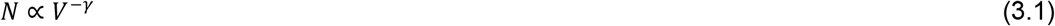

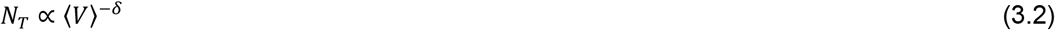

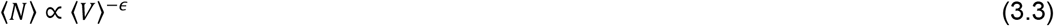

Combining Eqs. 3.1 with 2.1 and 3.2 with 2.2, and 2.2 with 3.3 imply that

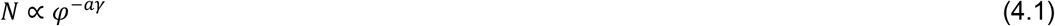

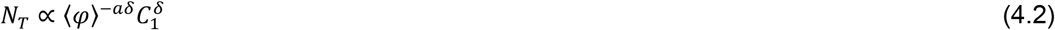

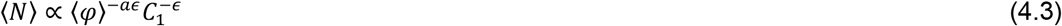

Secondly, we will use these derived predictions to, on the one hand, simplify metagenomic shotgun sequencing theory, and on the other, to make new predictions of metabolic ecology as applied to metagenomics, including the temperature dependence of metagenomic structure (abundance and richness of genes and groups of genes, i.e., families, metabolisms). In metagenomics, the abundance of sequenced reads of a population and gene can be predicted from DNA sequencing theory. The number of sequence reads of individuals of a taxon (*R*_P_) depends on its genome size (*φ*) and abundance (*N*) and mean genome size of the community (Beszteri et al. 2010, Bewick et al. 2017) as

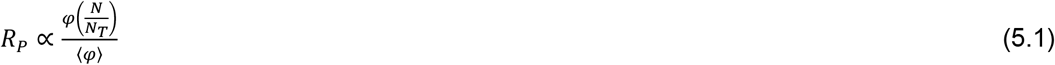

where *R*_P_ is the number of reads (reads of individuals), *φ* is genome size in (bp), *N* is population proportional abundance (individuals/total number of individuals, and ⟨*φ*⟩ is the mean genome size of the community (bp/total number of individuals). Replacing Eqs. 4.1 and 4.2 in eq. 5.1 implies that

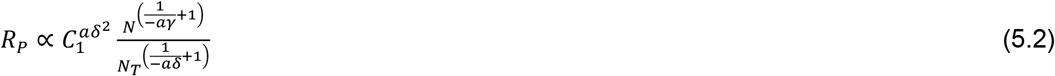

Similarly, despising duplicated gene copies (Treangen and Rocha), the number of reads of a gene sampled in a community (*R*_G_) depends proportionally on gene length (*L*) (bp), mean number of copies ⟨*n*⟩ (copies/individuals and inversely proportional to mean genome size (Beszteri et al. 2010) (bp/individuals) as

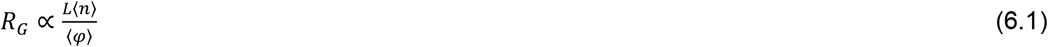

Where *R*_G_ (reads of gene copies), *L* (bp), ⟨*n*⟩ (copies/individuals), and ⟨*φ*⟩ is the mean genome size of the community (bp/individuals). Simplifying the units in the right side, [(bp)(copies/individuals)/(bp/individuals)], it can be seen that *R*_G_ measures in the number of copies of a gene. The number of (recently duplicated) gene copies scales with genome size (in microorganisms; Martínez-Núñez et al. 2013), which can be described by a power-law, *n* ∝ *φ*^*λ*^. Averaging we have

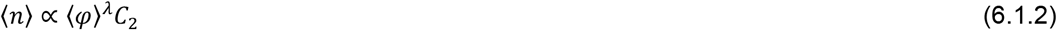

Where *C*_2_is a constant (see Savage 2004). But considering that 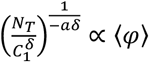 (Eq. 4.2), and combining Eq. 6.1.2 with 4.2 we have

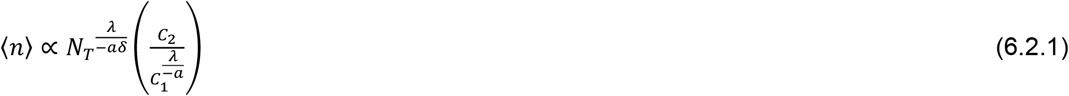

Now replacing 4.2 and 6.2.1 in 6.1 and assuming that the gene length (L) is approximately constant among samples (i.e. changes due to insertions and deletions do not imply major differences in the length of a particular gene across spatial gradients) gives

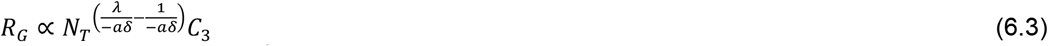

Where 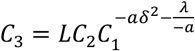

### Predicting the taxonomic community structure

We will integrate the volume-temperature rule with the abundance-size relationship to predict the dependence of population and assemblage abundance on temperature. It has been theoretically shown that size in ectotherms depends on temperature, due to their effects on growth and development (Zuo et al. 2012). The intraspecific volume-temperature rule can be approximated as

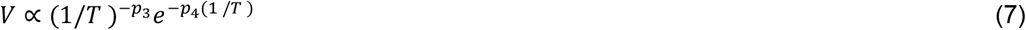

This prediction has been supported by data at the level of population in bacteria, archaea, and unicellular eukaryotes (Shehata and Marr 1975, Hjort and Bernander 1999, Montagnes and Franklin 2001, Gervais et al. 2003). Combining eq. 7 and eq. 3.1 implies

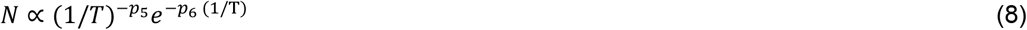

Where *p*_5_ = *p*_3_*γ*, and *p*_6_ = *p*_4_*γ*.

We can derive the temperature dependence of assemblage abundance by applying summation to Eq. 8 across taxa and assuming that temperature is the same and constant within (ectotherm) taxa in the community, resulting in

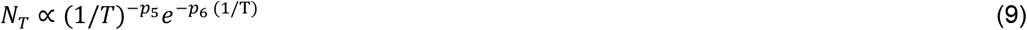

Let’s consider that the number of taxa (S; richness) scales with the number of individuals of an assemblage (e.g., Locey and Lennon 2016) according to

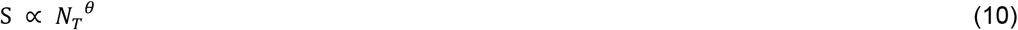

Now we reconcile the simplified DNA shotgun equation. 5.2 with predictions of metabolic ecology, Eqs. 8-10. Replacing 8 and 9 in 5.2 implies that the number of reads (of individuals) of a population depends on temperature

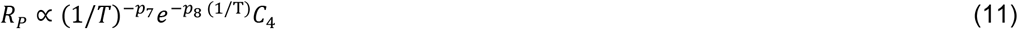

Where 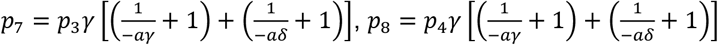, and 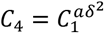.

The number of reads of groups of populations (*R*_T_) is the sum of the reads of all populations, i.e. *R*_T_ = Σ*R*_P_. To derive the temperature dependence of *R*_T_ we replace 8 and 9 in 5.2 and then apply summation, and as the temperature is a constant, as is the same in the community implies

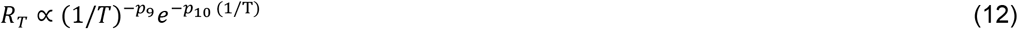

Where 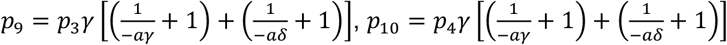.

### Predictions for the structure of metagenomes

We can simply assume a relationship between i) the number of genomes and the abundance of a gene in a population, ii) the summed number of genomes and the summed abundance of a gene in a community, and iii) the individual and the summed number of genomes in a community. Then, combining with equation 6.3 it is predicted eq. 13. Alternatively, we can directly replace 9 in 6.3, which predicts that the number of reads associated with a particular gene or the number of copies of a gene in an assemblage depends on temperature

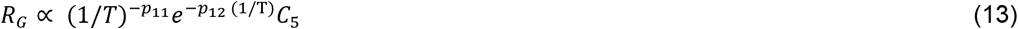

Where 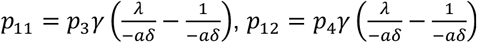, and 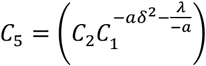.

Using this last derivation, we can predict the temperature dependence of taxa richness in a metagenomic sample. Combining eq. 6.3 and eq. 10 we have 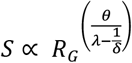 and then combining this expression with Eq. 13, we have

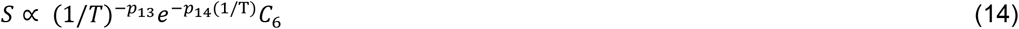

Where 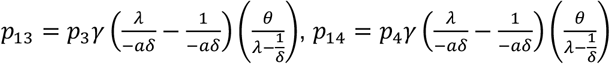, and 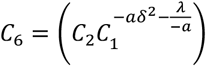.

The number reads of copies of genes of a group of related genes (*R*_M_; pathway, family) is the sum of the reads of copies *R*_M_ = Σ *R*_G_. We can derive the temperature dependence of *R*_M_by taking the summation in Eq. 13, which implies

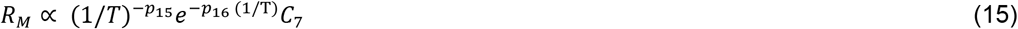

Where 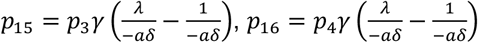, and 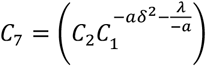.

To derive a prediction for the temperature dependence of the number of genes and groups of genes, we require three assumptions. i) a scaling among the mean genome size of a population 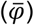 with the mean number of genes of a population (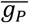; a relationship whose extent could vary among taxonomic domains but can be described as a simple power-law (Hou and Lin 2009);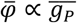, ii) after taking the summation of this previous relationship, a scaling among the whole genome size of a community 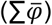 and the number of genes in a community 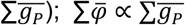, iii) after applying summation to equation 4.1 we have our third assumption, which is a scaling of the summed genome size 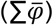 and the summed population abundance (i.e., community abundance; 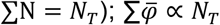. Combining these assumptions, we can further assume that the number (richness) of different genes (*g*_*C*_) and number (richness) of groups of genes (*g*_*M*_) depends mainly on the total number of individuals in the assemblage under analysis, thus:

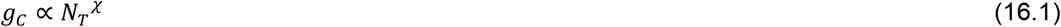

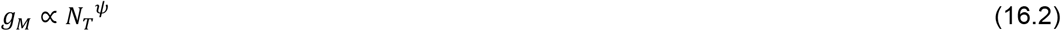

Combining 16.1 with 6.3 and then combining with Eq. 13, and combining 16.2 with 6.3 and then combining with Eq. 13 it is deduced that

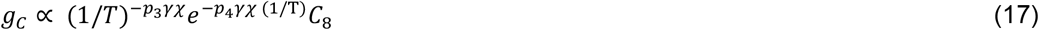

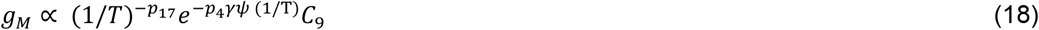

Where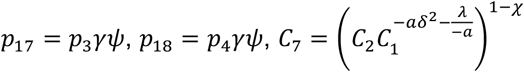, and 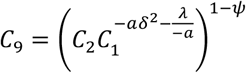.

### Testing predictions

We fitted our derived equations in a logarithmic form, using simple linear (for equations 4.1-4.2) or multiple linear regression (fit equations 11-15, 17, 18), to data collected from published studies. Eq. 4.1-4.2 are supported by data from hot springs communities (Arroyo et al. *In preparation*) and eq. 4.3 by data from human gut microbiome (Manor and Borenstein 2015), in all three cases showing a scaling pattern with a negative slope in a log scale (Figure 1). The predictions associated to equations 8, 9 and 14 concerning the role of temperature in affecting population abundance, assemblage abundance, and richness are similarly supported by the data (Figure 2). To fit the relationships between population and community abundance with temperature, we used as an example data from the Red Sea microbiome from the genus *Acholeplasma*, for any particular reason, but just because as fits well with the model. Both the population abundance of an *Acholeplasma palmae* and the abundance of the assemblage composed of three species of *Acholeplasma* varied non-linearly with temperature (Figure 2a and b, respectively). Finally, to fit predictions for taxonomic and functional richness, we used data from the Red Sea and global ocean microbiomes. The number of taxa varies non-linearly with temperature in the ocean microbiome for the fraction 0.22-3 in surface waters (SRF); Figure 2c. The predictions associated with equations 13 and 15 on the role of temperature on the abundance of genes and groups of genes in an assemblage were fitted using the abundance of the gene phosphogluconate dehydrogenase and the group of genes associated with the metabolism of photosynthesis (Fig. 3a-b). Finally, we fitted predictions deriving from equations 17-18 and associated with the role of temperature on the richness of genes and groups of genes in an assemblage. As an example, we used COG genes in the global ocean in negative latitudes, fraction 0.22-3 in the mesopelagic zone (MES; Fig. 3c), and KEGG modules in the global ocean (size fractions 0.22-1.6; Fig. 3d), both responding predictably to temperature.

**Fig. 1.**
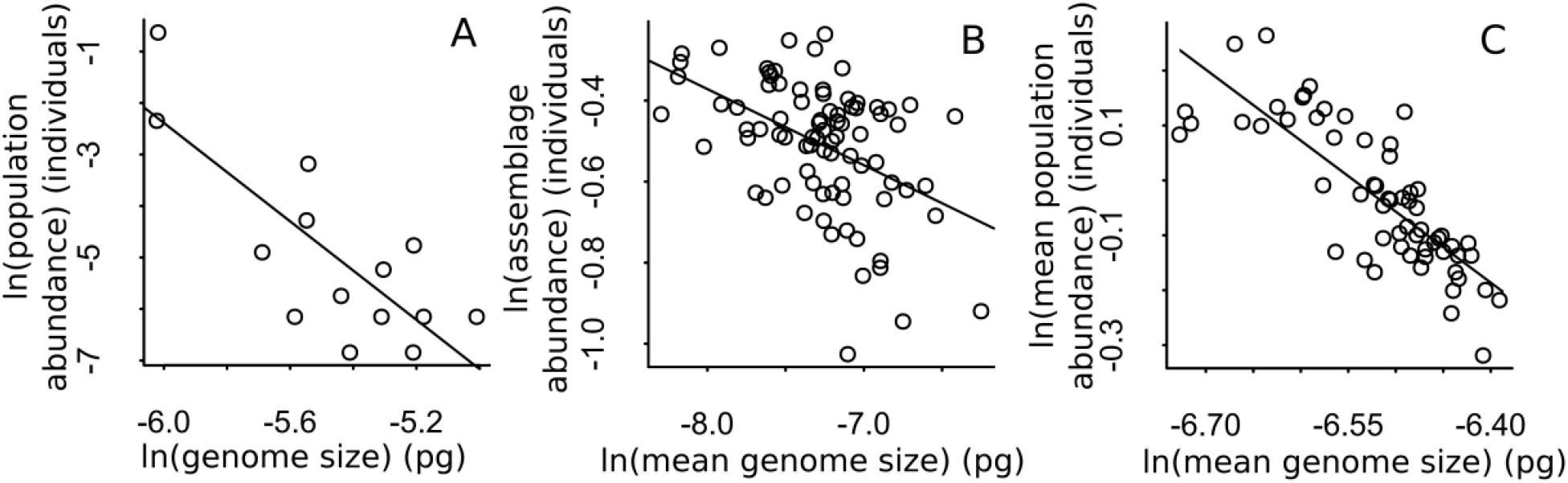
Scaling patterns of abundance (individuals) and genome size (picograms, pg). (a) local population abundance and reference genome size in a Patagonian hot spring bacterial mat, (b) assemblage abundance and mean genome size among 86 Patagonian hot spring bacterial mat communities, (c) global mean population abundance and mean cell volume in human gut.

**Fig. 2.**
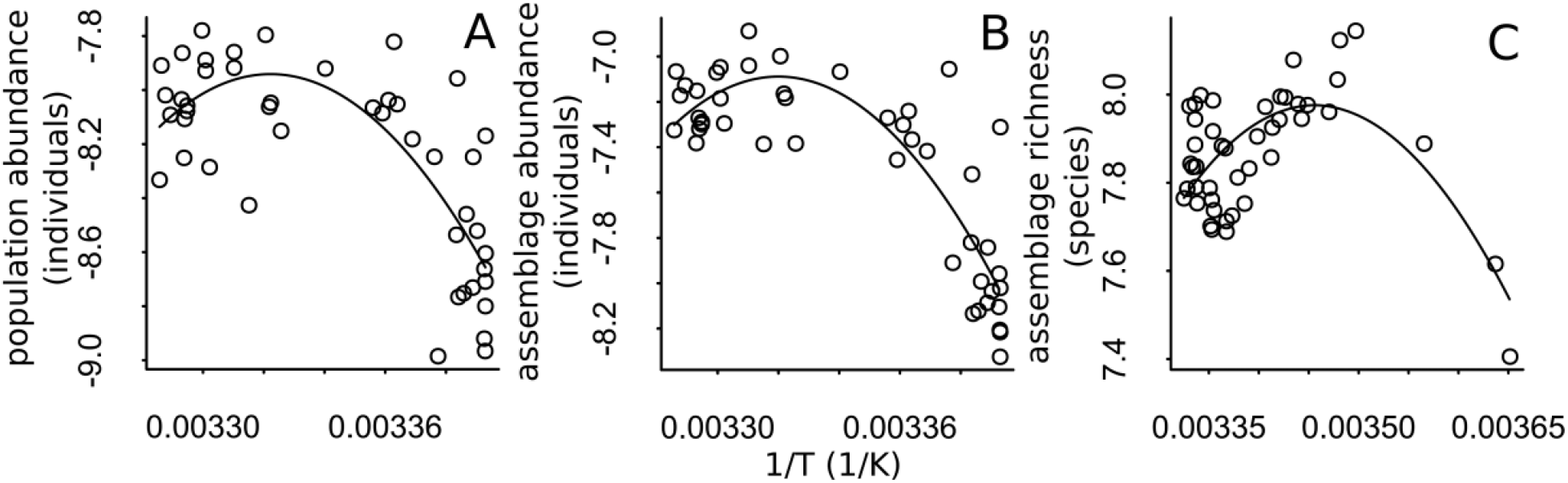
Temperature dependence of (a) ln(population abundance) (individuals) of *Acholeplasma palmae*, (b)ln (community abundance) (individuals) of *Acheoleplasma*, and (c) ln(taxa richness) (species) in the ocean microbiome in the fraction 0.22-3 in surface waters (SRF).

**Fig. 3.**
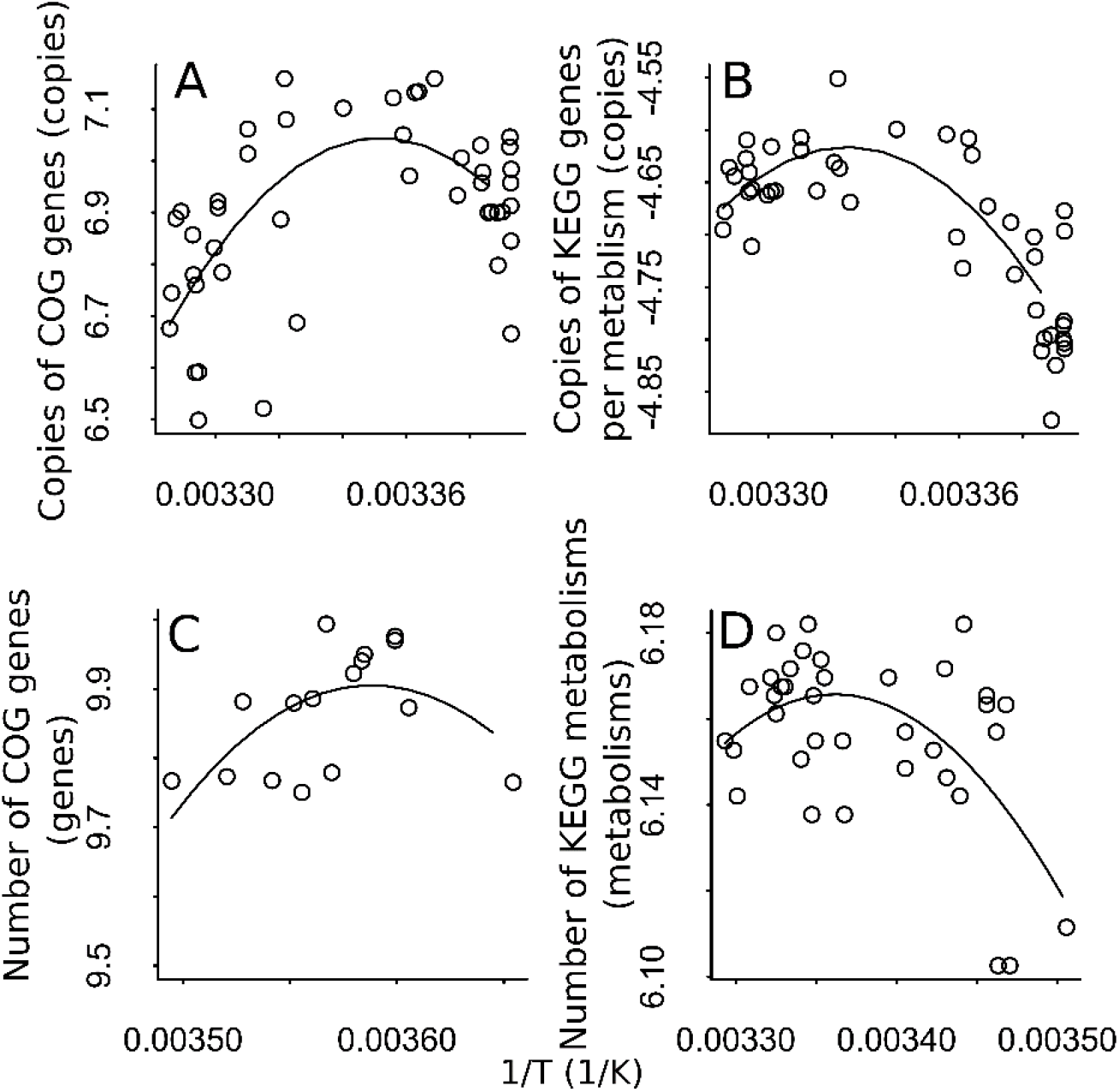
(a) ln(gene abundance) (copies) of the methylenetetrahydrofolate reductase NADPH (K00297) gene and temperature in the Red Sea, (b) ln(metabolism abundance) (copies) of bacterial secretion systems (Ko03070) genes and temperature in the Red Sea, (c) ln (COG genes) (genes) (0.22-3 mesopelagic zone-MES-in negative latitudes) in the global ocean, and (d) ln(KEGG modules) (groups of genes), particularly of size fraction0.22-1.6, in the global ocean and temperature.

## Discussion

An emergent challenge is to integrate theory with omic approaches (meta-proteomics, metatranscriptomics, metagenomics, etc.), which allows the study of the ‘functional’ basis of ecological patterns (Barberán et al. 2014). Meta-omic approaches started descriptively, but recently, theoretical studies have been done. For instance, biogeochemical models have been used to predict functions (Reed et al. 2014, Coles et al. 2017), and inversely, metagenomes have been used to predict environmental parameters (Gianoulis et al. 2009). In addition to geochemical substrates, temperature is often the most explicative irrespective of scale, environment or taxa, then we focused on this parameter. Among ecological theories, the metabolic ecological theory provides a mechanistic basis for its effects through different levels of organization. Here we took a deductive, static, deterministic approach to derive the mechanistic basis of ecological patterns found in microbiomes, particularly their genes and groups of genes, which are relevant as they perform ecosystem processes. We attempted to made three major contributions, i) to simplify DNA sequencing theory for metagenomics using metabolic ecology rules, particularly that the number of reads of a population depend only on population and assemblage abundance and the abundance of a gene depend only on total abundance, ii) Then we predicted the temperature dependence of community structure (i.e. abundance and richness of populations and assemblage), and iii) derived the temperature dependence of the abundance and richness of genes and groups of genes (i.e. families, pathways, etc). Below, we discuss relevant points.

First, we combined the relationships population abundance, cell size, and cell volume-genome volume to derive the relationship population abundance-genome size (Eq. 4.1), which was well supported by data (Fig. 1a). The mechanism and degree of dependence (slope of the relationships in log scale) underlying the relationship between genome size and cell size may be different for prokaryotes and unicellular eukaryotes (Cavalier-Smith 2005, Marshall et al. 2012, Mueller 2015, Gallet et al. 2017, Fernandez-de-Cossio-Diaz and Vazquez 2018). In both cases, it has been argued that cell size depends on genome size due to constraints on optimization of cellular physiological rates (metabolic, growth; West et al. 1999, Kozłowski et al. 2003, Shestopaloff 2016, Gallet et al. 2017). On the other hand the relationship between population abundance and body size is based on the hypothesis that the maximum number of individuals of a population (Nmax) a given ecosystem can sustain depends on the ratio of available resources (R) and the rate at which these individuals exploit those resources (B; N~R/Nmax; Enquist et al. 1998). Using these derived relationships for population abundance and genome size and assemblage abundance and mean genome size (Eqs. 4.1 and 4.2, respectively), we simplified DNA sequencing model of Beszteri et al. (2010). The model by Beszteri et al. (2010) was used to develop methods to estimate taxa abundance in metagenomic samples (Nayfach and Pollard 2015), however until now it had not been considered that due to the scaling rules between abundance, cell volume and genomic mass the number of reads is proportional to abundance, and therefore reads can be used directly to compare community structure across environmental gradients.

Second, we derive relationships for community structure and its dependence on temperature (Eqs. 11-14) and show examples (Fig. 2) using DNA sequencing theory. We believed could be useful for mechanistic understanding to derive the temperature dependence of community structure as many metagenomic studies have shown that the relative or proportional abundance (of reads) of individuals and richness of microbial taxa change with temperature but, to the best of our knowledge, infrequently the underlying causes of this variation are discussed (e.g. Sunagawa et al. 2015, Menzel et al. 2015, Ding et al. 2015). According to our derivations, the abundance of an assemblage determines its richness, and both are determined by the distribution of the cell volume, which in turn is also determined by the temperature. It is worth mentioning that in metabolic ecology, predictions had already been derived for the carrying capacity (i.e., maximum population abundance; Savage et al. 2004), abundance of heterotrophic assemblages (Allen et al. 2007), and species richness (Allen et al. 2002, Allen et al. 2006, Allen et al. 2007). In the case of the derivation for carrying capacity, it was done using the temperature dependence of the growth rate, and in the case of the abundance of a local assembly (of heterotrophs), using the net primary productivity (Allen et al. 2007). On the other hand, species richness has been derived using different approaches, including population metabolism (Allen et al. 2002) and speciation rate (Allen et al. 2006).

Third, to derive Eqs. 17-18 (Fig. 3b-d), the underlying assumption is that, even considering duplicated and deleted genes, we can assume a scaling relationship between the abundance of groups of genes and the number (richness) of genes with assemblage abundance (i.e., the number of individuals in the assemblage). From a mechanistic point of view, this scaling is expected based on the population genetics theory, whereby polymorphism for a neutral locus is expected to increase with effective population size. Furthermore, since we sample a specific time and space of the community, for practical purposes, we can disregard other rates such as migration, or the emergence of new genes by duplication or deletion. This point of view is not new in ecology and has already been used not only for the scaling relationship between species and abundance (MacArthur and Wilson 1967, Hubbell 2001), and gene flow (Vellend 2003).

It is also important to mention that our theoretical framework corresponds to general deductions and not specific predictions that include expected values of parameters. This is because, although in some cases, such as for the abundance-volume relationship, there is a predicted value for the power of volume (−3/4; Enquist et al. 1998), for the rest of the equations, there is as yet no theory that makes specific predictions about the value of the parameters. However, the value of the parameters in Eq. 1.2 varies greatly, and can be predicted by i) clearing the parameter in the equation. 1.2, ii) now considering k as a variable and the rest of the terms as new parameters, and iii) replacing k with a variable that also predicts it, preferably using an equation also derived from a theory. For example, in chemical kinetic theory, the rate of reaction is not only dependent on temperature but also on pH, so, for example, if we have temperature response curves each at a different pH, we could also fit a model for the thermodynamic parameters. This approach has been taken previously to predict, for example, the activation energy (E) of the Arrhenius equation in the relation number of species and temperature using area, which is another variable that also predicts the number of species (Wang et al. 2009). For example, in the case of genomic community structure, any of the predicted variables depends on the number of total reads (Tringue et al. 2005), so this variable could predict the value of the parameters. In particular, it could be predicted, for example, that the value of parameters would vary predictably from genes to metabolisms and species.

Regarding the fit of our compiled data, we focused on showing the predictability of patterns, and for simplicity, we showed a single example; nonetheless, in recent studies, it is possible to find more examples (see Ding et al. 2015, supplementary material of Sunagawa et al. 2015; Thompson et al. 2016 Fig S5 Table S1). In general our fits (Table 1) had an intermediate level of explained variance, which could be related to the stochasticity of the ocean environment but also due to other variables that can also be important predictors the abundance and richness of genes of populations and communities such as nutrients (Mende et al. 2017), and other physical and chemical variables that also affect enzyme reactions rates such as pH and salinity (Lozupone and Knight 2007, Fierer and Jackson 2016).

**Table 1.**
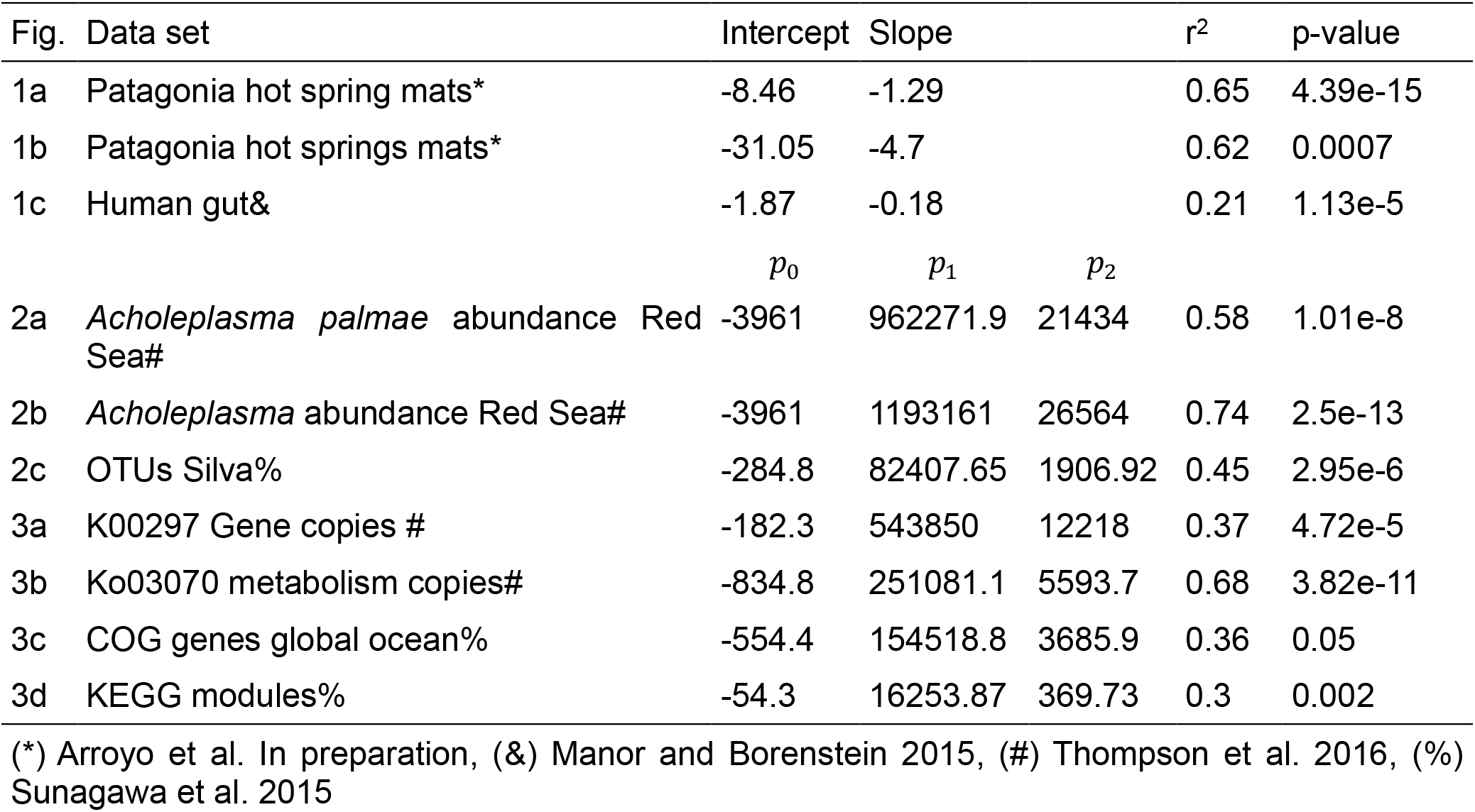
Estimated parameters of and goodness of fit of Figs. 1–3.

We visualize many other applications of metabolic theory of ecology on metagenomics, to study, for instance, functional redundancy (e.g. measured for instance as the quotient of taxa over functions) should also depend on temperature, ii) further developments towards theoretical metagenomics could include predictions of gene beta diversity, and iii) population genomic diversity could be studied using genome-centric metagenomics or single-cell genomics (Swan et al. 2013). Moreover, further studies can now extend this approach to empirical relationships often studied in metabolic ecology and macroecology, such as the study of the distribution of gene abundances, distribution of gene and species richness, distribution of genome sizes scaling relationships of gene diversity, the effects of stoichiometry on gene richness (se for reference Allen and Gillooly 2009), among others. Our predictions are also applicable to the microbiomes of host-associated ecto- and endotherms, which have a range in their body temperatures spanning 10°C (Clarke and Rothery 2008). For instance, in ectotherms, a few recent studies using amplicon sequencing have shown that the diversity of microbial communities changes with temperature (Kohl and Yahn 2016, Bestion et al. 2017, Carrell et al. 2017).

In summary, by integrating the first principles of metabolic ecology and high-throughput sequencing theories, we derived predictions for the structure of microbial community genomes, representing a step forward over comparative approaches. Predictions regarding these properties were supported by global microbiome data from human, ocean, and terrestrial microbiomes. We visualize a great potential in integrating metagenomics with ecological theories, which could lead to a better understanding of the mechanism behind the observed patterns, together with specific testable predictions that could serve to easily anticipate phenomena and take informed decisions for management, especially in problems involving ecosystem processes.

## Methods

We used a deductive logical approach to derive predictions (Gunawardena 2014, Kirchner and Kirchner 2014) to integrate some of the principles of three theories: metabolic ecology, population genomics, and DNA sequencing theory. Theoretical predictions were performed by making transitive relationships between two relationships to derive a third. Each relationship had a hypothesized causal explanation grounded on basic ecological principles, making the transitive deductions also mechanistic.

To fit the models to data, information was obtained directly from tables of published articles, from the supplementary data, or extracted from figures using a plot digitizer. Particularly, we used data from a study of bacteria in Patagonia hot springs (Arroyo et al., In preparation), the human microbiome (Manor and Borenstein 2015) to test the relationships between abundance and genome size, Red Sea (Thompson et al. 2016) and global ocean microbiome (KEGG genes and modules) (Sunagawa et al. 2015) to test the relationship between community structure of species, genes, and metabolisms with temperature. The relationship between abundances and genome sizes was fitted using linear regression to a power-law on a logarithmic scale, *Y* = *aX*^*b*^, ln *Y* = ln *a* + *b* ln *X*. The relationships of temperature dependence were fitted using multiple linear regressions to a power law with exponential cut-off on a logarithmic scale: *Y* = *aX*^*b*^*e*^*cX*^, In *Y* = In *a* + *b* ln *X* + *cX*. Regressions were implemented in the language R (R Core Team 2012). The goodness of fit and significance were assessed based on R^2^ and P-value, respectively.

## Acknowledgments

JIA acknowledges support from a CONICYT National Doctoral Scholarship N° 21130515

