## Supplementary Material for "Metabolic ecology, microbial community structure, and gene-centric metagenomics"

Tabla S1. List of variables/constants, their symbols, and units.

| Variable/constant | Symbol | Unit |
|---|---|---|
| (Enzyme) reaction rate constant (first-order reaction) | k | 1/s |
| Temperature | T | K |
| Boltzmann constant | $k_B$ | $(1.38*10^{-23})$ J/K |
| Planck constant | h | $(6.62 * 10^{-34})$ J/s |
| Gas constant | R | (8.31) J/Kmol |
| Free enthalpy-change or Gibbs energy | ΔG | J |
| Enthalpy change | ΔH | J |
| Heat capacity change | ΔC | J/K |
| Entropy change | ΔS | J/K |
| Entropy change at $T_0$ | $\Delta S_{T0}$ | J/K |
| A reference initial temperature, often 25°C | $T_0$ | K |
| Population abundance | N | individuals/área or $uL^3$ |
| Cell volume | V | $\mu m^3$ |
| Number of species of a community | S | species |
| Number of genes of a population | $g_P$ | genes |
| Number of genes of a community | $g_C$ | genes |
| Number of group of genes of a community | $g_M$ | groups of genes |
| Abundance of an 'assemblage' | $N_T$ | individuals/$uL^3$ |
| Genome size | φ | bp or pg |
| Gene length | L | bp |
| Copy number | n | copies |
| Number of reads of a species | $R_P$ | individuals |
| Number of reads of a gene | $R_G$ | copies |
| Number of reads of a group of genes | $R_M$ | copies |

*Selection coefficient can be measured a quotient of the relative fitness of two genotypes therefore is dimensionless
